# Sexually dimorphic effects of pexidartinib on nerve injury-induced neuropathic pain in mice

**DOI:** 10.1101/2023.10.10.561386

**Authors:** Fumihiro Saika, Yohji Fukazawa, Yu Hatano, Shiroh Kishioka, Shinjiro Hino, Kentaro Suzuki, Norikazu Kiguchi

**Author notes:** Corresponding author: Norikazu Kiguchi, Ph.D.

## Abstract

**Background:** It is well-established that spinal microglia and peripheral macrophages play critical roles in the etiology of neuropathic pain; however, growing evidence suggests sex differences in pain hypersensitivity owing to microglia and macrophages. Therefore, it is crucial to understand sex- and androgen-dependent characteristics of pain-related myeloid cells in mice with nerve injury-induced neuropathic pain.

**Methods:** The current study was performed using normal male and female mice, as well as gonadectomized (GDX) male mice. To deplete microglia and macrophages, pexidartinib (PLX3397), an inhibitor of the colony-stimulating factor 1 receptor, was orally administered, and mice were subjected to partial sciatic nerve ligation (PSL). Immunohistochemistry was performed to visualize microglia and macrophages, and PSL-induced mechanical allodynia was evaluated using the von Frey test.

**Results:** Following PSL induction, healthy male and female mice and male GDX mice exhibited similar levels of spinal microglial activation, peripheral macrophage accumulation, and mechanical allodynia. Treatment with PLX3397 significantly suppressed mechanical allodynia in normal males; this was not observed in female and GDX male mice. Sex- and androgen-dependent differences in the PLX3397-mediated preventive effects were observed on spinal microglia and dorsal root ganglia (DRG) macrophages, as well as in expression patterns of pain-related inflammatory mediators in these cells. Conversely, no sex- or androgen-dependent differences were detected in sciatic nerve macrophages, and inhibition of peripheral CC-chemokine receptor 5 prevented neuropathic pain in both sexes.

**Conclusion:** Collectively, these findings demonstrate the presence of considerable sex- and androgen-dependent differences in the etiology of neuropathic pain in spinal microglia and DRG macrophages but not in sciatic nerve macrophages. Given that the mechanisms of neuropathic pain may differ among experimental models and clinical conditions, accumulating several lines of evidence is crucial to comprehensively clarifying the sex-dependent regulatory mechanisms of pain.

## Introduction

Neuropathic pain attributed to nerve injury and disease is a prevalent problem that not only worsens the patient’s quality of life but also represents an enormous economic burden globally [1, 2]. Given that current treatments are often ineffective in treating the symptoms of neuropathic pain, such as allodynia, identifying the etiology of neuropathic pain is imperative for developing novel therapeutics. Several lines of evidence suggest that the interaction between neurons and non-neuronal cells through growth factors and inflammatory mediators is responsible for chronic neuroinflammation in the peripheral nervous system (PNS) and central nervous systems (CNS) [3, 4]. Therefore, the characterization of non-neuronal cells that largely contribute to neuropathic pain in different experimental models has received considerable attention.

At the site of nerve injury and in the dorsal root ganglia (DRG), macrophages are increased and activated in response to chemokines and hematopoietic growth factors produced by glial cells, such as Schwann cells, and the cell bodies of sensory neurons [5–7]. Among diverse macrophage subsets, inflammatory macrophages induced by colony-stimulating factor 1 (CSF1) can produce various inflammatory mediators (e.g., interleukin [IL]-1β and CC-chemokines) that alter the sensory transmission in the axons and cell bodies of primary sensory neurons, eliciting abnormal reactivity to painful or innoxious stimuli (peripheral sensitization) [8, 9]. Accordingly, the inhibition of inflammatory macrophages or macrophage-derived painful mediators could successfully attenuate hypersensitivity in different models of neuropathic pain, indicating the critical role of peripheral macrophages in the pathophysiology of neuropathic pain [10, 11].

Typically, microglia play a crucial role in maintaining homeostasis in the CNS, whereas reactive microglia contribute to several chronic diseases associated with neuroinflammation [12, 13]. Following peripheral nerve injury, upregulated CSF1 is released from the terminals of damaged sensory neurons in the spinal dorsal horn (SDH). Similar to peripheral macrophages, CSF1 induces the proliferation and morphological activation of spinal microglia [7, 14], and various soluble factors, including inflammatory cytokines, growth factors, and lipid mediators, participate in the hyperexcitation of pain-processing neurons in the SDH (central sensitization) [15, 16]. The administration of microglial inhibitors has been shown to substantially alleviate hypersensitivity, whereas selective microglial activation elicits pain behaviors [17, 18]. These findings suggest that the activation of spinal microglial is a key mechanism underlying neuropathic pain. However, accumulating evidence suggests the presence of sex differences in the role of spinal microglia in neuropathic pain. For example, the administration of minocycline, a typical inhibitor of microglia, and inhibitors of key signaling molecules for reactive microglia or depletion of microglia were shown to suppress neuropathic pain in male but not female mice [19, 20]. Moreover, we have previously demonstrated that chemogenetic manipulation of spinal microglia could control neuropathic pain in male but not female mice [21], indicating the sex-dependent contribution of spinal microglia to neuropathic pain. These microglial characteristics have been defined by transcriptomic analyses revealing sex-dependent inflammatory profiles [22, 23]. Considering neuropathic pain, sex-dependent differences in peripheral macrophages remain poorly established when compared with those in spinal microglia. Nevertheless, in some cases, male macrophages are likely related to pain hypersensitivity [24, 25].

Given the presence of notable sex differences in the etiology of neuropathic pain in spinal microglia (and peripheral macrophages) [26, 27], it is pivotal to identify critical regulators that determine the sex-dependent characteristics of pain-related myeloid cells in both the PNS and CNS. As CSF1 is an essential factor involved in the proliferation and activation of macrophages and microglia [28], we evaluated the inhibitory effects of systemic exposure to pexidartinib (also called PLX3397), a CSF1 receptor (CSF1R) inhibitor [29], on nerve injury-induced neuropathic pain in mice. Importantly, we verified that androgen levels determine the effectiveness of PLX3397 and sex-dependent characteristics of macrophages and microglia underlying neuropathic pain.

## Methods

### Mice

All animal experiments were approved by the Animal Research Committee of Wakayama Medical University and were performed in accordance with the in-house guidelines for the care and use of laboratory animals of Wakayama Medical University and the Animal Research: Reporting of In Vivo Experiments (ARRIVE) guidelines. Male and female C57BL/6 mice (4–8 weeks old) were purchased from SLC (Hamamatsu, Japan) and used for each experiment at 8–12 weeks of age. All mice were housed in groups of 5–6 in plastic cages maintained under controlled temperature (23–24°C), humidity (60–70%), and a 12-h dark/light cycle, with free access to food and water.

### Partial sciatic nerve (SCN) ligation (PSL) model

Mice were subjected to PSL as previously described [30, 31]. Briefly, under isoflurane anesthesia, the left common SCN of each mouse was exposed at the mid-thigh level by performing a small skin incision on one side, hereafter referred to as the ipsilateral side. Approximately one-third of the SCN thickness was tightly ligated with a silk suture (Natsume Seisakusho, Tokyo, Japan), followed by suturing of muscle and skin layers and sterilization of the surgical area with povidone–iodine. The untreated right limb was considered the contralateral limb.

### Administration of PLX3397

To achieve macrophage and microglia depletion *in vivo*, PLX3397 (MedChemExpress, Monmouth Junction, NJ, USA), a CSF1R inhibitor, was formulated into the AIN-76A rodent diet (Research Diets, New Brunswick, NJ, USA) at 290 mg/kg. The PLX3397 dose was established based on a previous report [29]. Mice had free access to the PLX3397-formulated diet for 1–3 weeks instead of normal food, as PLX3397 is orally active. The AIN-76A rodent diet was used as the control.

### Reverse transcription-quantitative polymerase chain reaction (RT-qPCR)

Mice were euthanized using isoflurane, and fresh dorsal horns of the lumbar (L4-5) SDH, DRG, and SCN samples were collected in RNAlater solution (Thermo Fisher Scientific, Waltham, MA, USA). Total RNA was isolated from the tissues using the TRIzol Plus RNA Purification Kit (Thermo Fisher Scientific) following the manufacturer’s instructions. Briefly, tissues were placed in a 1.5-mL RNase-free tube and homogenized with TRIzol reagent. Chloroform was added to each sample, which was then centrifuged at 4°C for 15 min. The aqueous phase containing RNA was transferred to a fresh tube, and RNA was isolated using a purification column. Total RNA extract (1 µg) was incubated with random primers (Promega, Madison, WI, USA) at 70°C for 5 min and subsequently cooled on ice. Samples were converted into cDNA by incubation with M-MLV Reverse Transcriptase (Promega) and dNTP Mix (Promega). qPCR was performed using an AriaMx Real-Time PCR System (Agilent Technologies, Santa Clara, CA, USA) with template cDNA (10 ng), primers for each gene (Thermo Fisher Scientific), and SYBR Premix Ex Taq II (Takara Bio Inc., Kusatsu, Japan). The primer sequences are listed in **Table S1**. Reactions were performed under the following conditions: 3 min at 95°C, followed by 45 cycles of step two comprising 10 s at 95°C and 30 s at 60°C. Fluorescence intensities were recorded, and data were normalized to β-actin (*ACTB*).

### Immunohistochemistry (IHC)

The lumbar (L4–5) spinal cord, DRG, and SCN were harvested from euthanized mice after transcardiac perfusion in phosphate-buffered saline (PBS) and fixed in 4% paraformaldehyde/phosphate buffer solution. Subsequently, specimens were post-fixed in 4% paraformaldehyde and cryoprotected in 30% sucrose/PBS solution at 4°C overnight. After embedding in a freezing compound (Sakura, Tokyo, Japan), frozen tissues were longitudinally cut into 30-μm thick (spinal cord) or 12-μm thick (DRG and SCN) sections using a cryostat (Leica Microsystems, Wetzlar, Germany) and floated in PBS (spinal cord) or mounted on a glass slide (DRG and SDN). The sections were treated with PBS containing 0.1 % Triton X-100 (PBST) for 1 h and then blocked with 5% donkey serum at 15–25°C for 2 h. Next, the sections were incubated with primary antibodies against ionized calcium-binding adapter molecule 1 (IBA1) (rabbit polyclonal, 1:1000; Wako, Japan), NeuN (mouse monoclonal, 1:500; Millipore, Billerica, MA, USA), and F4/80 (rat monoclonal, 1:200; Cederlane, Burlington, Canada) at 4°C overnight. Subsequently, the prepared sections were rinsed in PBST and incubated with fluorescence-conjugated secondary antibodies (1:200; Thermo Fisher Scientific) at 15–25°C for 2 h. Finally, the sections were washed with PBS, mounted on glass slides, and covered with a cover slip using Immunoselect Antifading Mounting Medium DAPI (Dianova, Hamburg, Germany). Fluorescent images were obtained using a confocal laser scanning microscope (Olympus, Tokyo, Japan). The number of IBA1^+^ cells within the lamina I-III of the SDH and the number of IBA1^+^ and F4/80^+^ macrophages within the DRG or SCN were measured in a square of 200 × 200 μm^2^ area using FLUOVIEW software.

### von Frey test

The 50 % paw-withdrawal threshold was determined using the von Frey test, as described previously [17]. Briefly, mice were individually placed on a metal mesh (5 × 5 mm) grid floor and covered with an acrylic box. After adaptation for 2-3 h, calibrated von Frey filaments (Neuroscience, Tokyo, Japan) were applied to the middle of the plantar surface of the hind paw through the bottom of the mesh floor. The filament set used in the current study consisted of nine calibrated von Frey filaments: 0.02, 0.04, 0.07, 0.16, 0.4, 0.6, 1.0, 1.4, and 2.0 g. In the up-down method, the test started with the application of 0.4 g filaments. Quick withdrawal, shaking, biting, or licking of the stimulated paw was considered a positive paw-withdrawal response. In the absence of a paw-withdrawal response to the selected force, the next stronger stimulus was applied. The next weaker stimulus was selected during paw withdrawal, in accordance with Chaplan’s procedure [32], upon crossing the response threshold for the first time (two responses straddling the threshold). Based on responses to the von Frey filament series, the 50 % paw-withdrawal threshold was calculated.

### Hargreaves test

The withdrawal threshold was determined using the Hargreaves test, as described previously [33]. Mice were placed on a glass sheet and covered with a clear acrylic box. After adaptation for 2–3 h, a radiant heat source (IITC 390 Plantar Test Analgesia Meter, Neuroscience) was placed under the glass sheet and applied to the plantar surfaces of both hind paws. Withdrawal latencies were evaluated based on the mean latency of three stimulations. A cut-off latency of 15 s was set to avoid tissue damage.

### Rotarod test

A rotarod apparatus (Panlab, Barcelona, Spain) was used to assess the motor function. A few days before the test, the mice underwent pre-training to ensure habituation to the rod rotating at various speeds (10 and 15 rpm). The mice were placed on the rod facing the direction opposite to the rotation and ambulated until they fell from the rod, or the maximum observation time had elapsed (the maximum observation time was 180 s). For all experiments, the latency and rotational velocity at which the animal fell from the rotarod were recorded. After training for a few days, mice rested for one day, followed by testing (three trials per day) on a rod rotating at various speeds (10 and 15 rpm), with 10-15 min intervals between each trial. The mean time spent on the rotarod apparatus was measured at each velocity [21].

### Gonadectomy

For orchiectomy, male mice were anesthetized with isoflurane. After the aseptic preparation of the surgical site, a small incision was made at the lower abdomen to exteriorize the testes, vas deferens, and spermatic blood vessels. Blood vessels and vas deferens were cauterized, and the testes were removed. The skin incision was closed with sutures, and the surgical area was sterilized with povidone–iodine. Four weeks post-surgery, the animals were used for the experiments.

### Serum testosterone measurement

Mice were euthanized by decapitation, and trunk blood was collected in a separate microtube (Sansho, Tokyo, Japan). In accordance with the manufacturer’s instructions, collected blood samples were maintained in the tube for 30 min, followed by centrifugation at 4°C for 5 min. The upper phase, containing the separated serum, was transferred to a fresh tube. Serum testosterone levels were measured using liquid chromatography-tandem mass spectrometry, which was performed by Asuka Pharmaceutical (Tokyo, Japan).

### Perineural administration

A CC-chemokine receptor 5 (CCR5) antagonist (maraviroc; Tocris Biosciences, Bristol, UK) was dissolved in dimethyl sulfoxide and diluted with sterile PBS for further use. A perineural injection was performed as described previously [31, 34]. Under isoflurane anesthesia, the agents (10 μL) were injected without a skin incision into the region surrounding the SCN using a microsyringe fitted with a 30-gauge needle connected to a cannula. After PSL induction, maraviroc injections were administered daily for seven consecutive days (days 0–6).

### Statistical analysis

Data are presented as the mean ± standard error of the mean (SEM). Statistical analyses were performed using the Student’s t-test, one-way analysis of variance (ANOVA) followed by Tukey’s multiple comparison test, or two-way ANOVA followed by Bonferroni’s multiple comparison test, as appropriate. Statistical significance was set at P < 0.05.

## Results

### Upregulation of microglial markers and inflammatory mediators in the SDH following PSL

Using RT-qPCR, we first investigated whether mRNA expression levels of microglial markers (IBA1, CD68, CX3C-chemokine receptor 1 [CX3CR1], and CSF1R), IL-1β, CC-chemokine ligand 3 (CCL3), and its receptor, CCR5, differed in the SDH of male and female mice following PSL induction. General microglial markers and inflammatory mediators were similarly upregulated in both male and female mice, and only IBA1, CSF1R, and CCR5 expression levels on day 7 were slightly higher in female mice than those in male mice. Expression levels of IL-1β and CCL3 in males showed similar expression patterns to those in females (**Fig. 1**).

**Figure 1.**
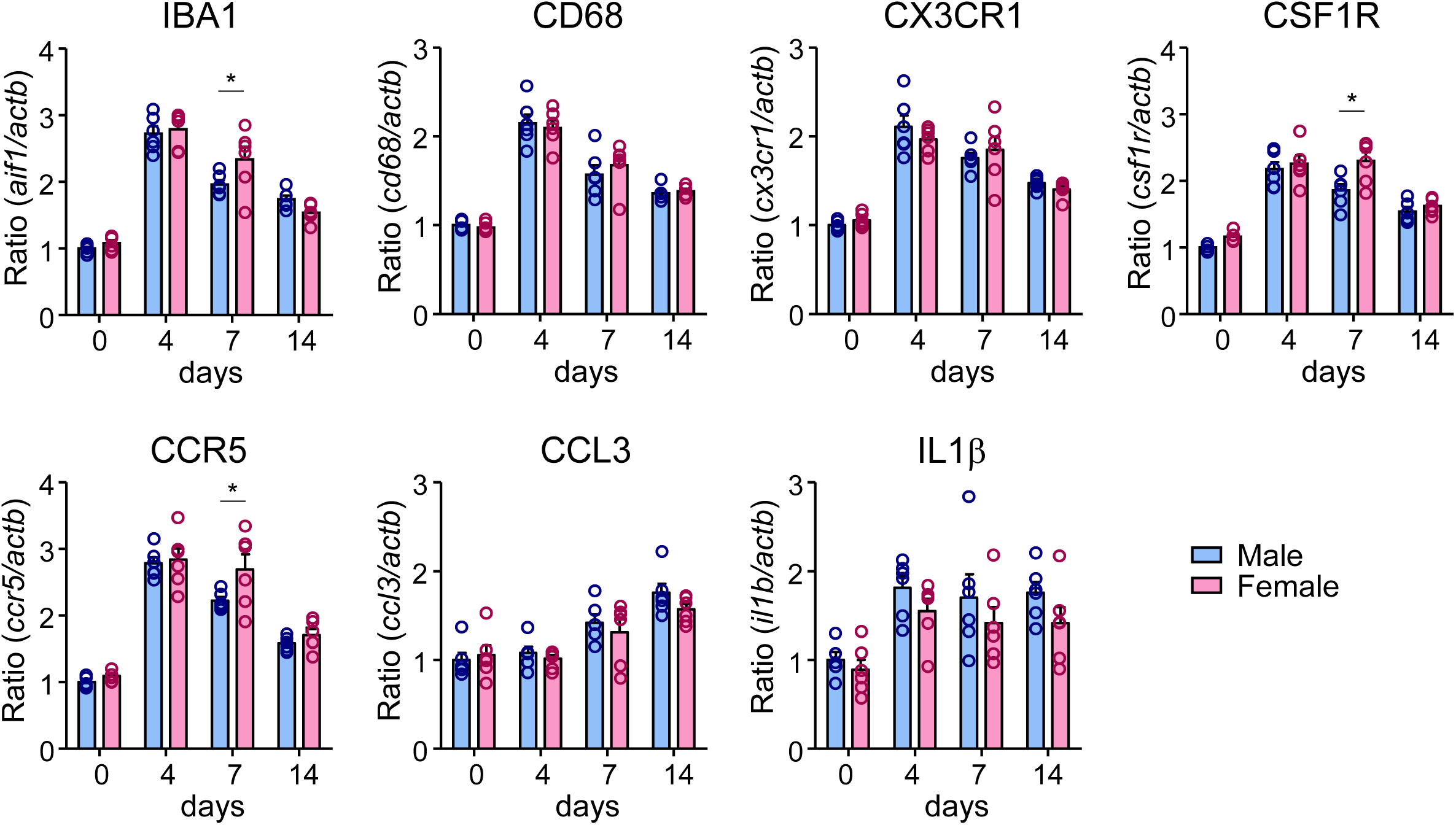
Upregulation of microglial markers and inflammatory mediators in the SDH following PSL. Male and female mice were subjected to PSL, and the lumbar SDH was collected. mRNA expression levels of *aif1*, *cd68*, *cx3cr1*, *csf1r*, *ccr5*, *ccl3*, and *il1b* on days 0 (pre), 4, 7, and 14 after PSL were analyzed by RT-qPCR. Data are presented as the mean ± standard error of the mean (SEM); n=5-6. \**P*<0.05 versus Male. PSL, partial sciatic nerve ligation; RT-qPCR, reverse transcription-quantitative PCR; SDH, spinal dorsal horn.

### Sexually dimorphic effects of PLX3397 on spinal microglial activation and mechanical allodynia following PSL

To determine the PLX3397-mediated effects on microglia and neuropathic pain, we used RT-qPCR to determine the time course of microglial ablation in the SDH following PLX3397 treatment. Following treatment with PLX3397, mRNA expression levels of IBA1 were significantly decreased at one-week post-treatment, decreasing further at two and three weeks. Treatment with PLX3397 did not affect the expression level of GFAP, an astrocytic marker, in the SDH at two weeks post-treatment (**Fig. S1A**). PLX3397 treatment did not affect the basal mechanical pain threshold, thermal withdrawal latency, or time on the rotarod (10 and 15 rpm) after two weeks of PLX3397 treatment (**Fig S1B-D**).

In male and female mice fed the control diet, the mechanical pain threshold was decreased on the ipsilateral side after PSL, indicating the development of mechanical allodynia. PSL-induced mechanical allodynia did not develop in PLX3397-treated male mice; however, PLX3397 treatment did not affect the mechanical pain threshold on the contralateral side. Conversely, PLX3397 treatment did not suppress PSL-induced mechanical allodynia in female mice, suggesting the sex-dependent effects of PLX3397 on neuropathic pain (**Fig. 2B**). According to IHC results, IBA1^+^ microglia were markedly increased on the ipsilateral side of the SDH on day 4 after PSL when compared with those on the contralateral side in both male and female mice. Treatment with PLX3397 markedly reduced the number of IBA1^+^ microglia on the ipsilateral and contralateral sides in both male and female mice, as confirmed by quantitative analysis (**Fig. 2C, D**). Notably, the number of remaining IBA1^+^ microglia in female mice was significantly greater than that in male mice, suggesting that male microglia are more susceptible to PLX3397.

**Figure 2.**
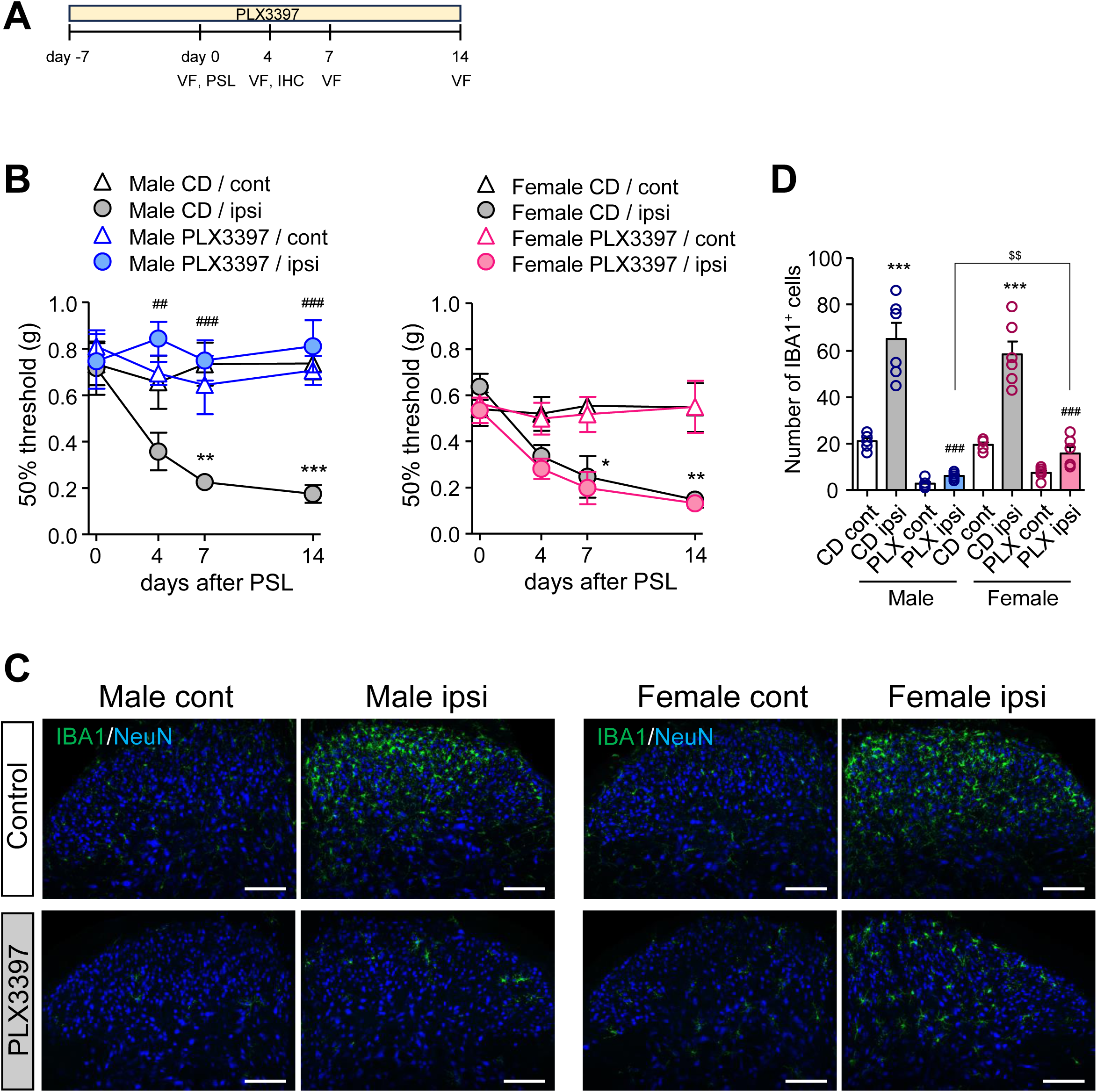
Sex-dependent PLX3397-mediated suppressive effects on mechanical allodynia and microglial activation in SDH following PSL. Male and female mice were fed a control diet (CD) or pexidartinib (PLX3397) diet and subsequently subjected to PSL. A) Time schedules of experiments. B) Effects of PLX3397 on the 50% mechanical threshold in male and female mice were assessed using the up-down method with the von Frey test. C) Effects of PLX3397 on the expression of IBA1 in the SDH of male and female mice were analyzed using IHC. Scale bars=100 μm. D) Quantitative analyses of the number of IBA1^+^ cells in the lamina I-III in the SDH on day 4 after PSL. Data are presented as the mean ± standard error of the mean (SEM); n=6. \*\*\**P*<0.001, \*\**P*<0.01, \**P*<0.05 versus CD cont. ^###^*P*<0.001, ^##^*P*<0.01 versus CD ipsi. ^$$^*P*<0.01 versus PLX ipsi. IHC, immunohistochemistry; PSL, partial sciatic nerve ligation; SDH, spinal dorsal horn.

### Roles of androgens on the suppressive effects of PLX3397 on spinal microglial activation and mechanical allodynia following PSL

Given that sex hormones, such as androgens, reportedly participate in pain regulatory mechanisms [35, 36], we examined whether androgen levels correlated with the sex-specific suppressive effects of PLX3397 on neuropathic pain. In gonadectomized (GDX) male mice, serum testosterone levels were below the detection limit at four weeks post-gonadectomy, whereas serum testosterone levels in females were significantly lower than those in males but were detectable (**Fig. 3B**). Next, we examined the effects of PLX3397 on microglia and neuropathic pain in GDX mice. Similar to female mice, PLX3397 treatment failed to suppress PSL-induced mechanical allodynia in male GDX mice, suggesting the androgen-dependent effects of PLX3397 on neuropathic pain (**Fig. 3C**). Based on IHC analysis, the ipsilateral side of the SDH showed a marked increase in IBA1^+^ microglia on day 4 after PSL when compared with the contralateral side in both normal and GDX male mice. In both normal and GDX male mice, treatment with PLX3397 dramatically reduced the number of IBA1^+^ microglia on the ipsilateral and contralateral sides (**Fig. 3D, E**). Consistent with the behavioral assay results, the number of remaining IBA1^+^ microglia in GDX male mice was significantly greater than that in normal male mice, suggesting that androgen levels determine susceptibility to PLX3397.

**Figure 3.**
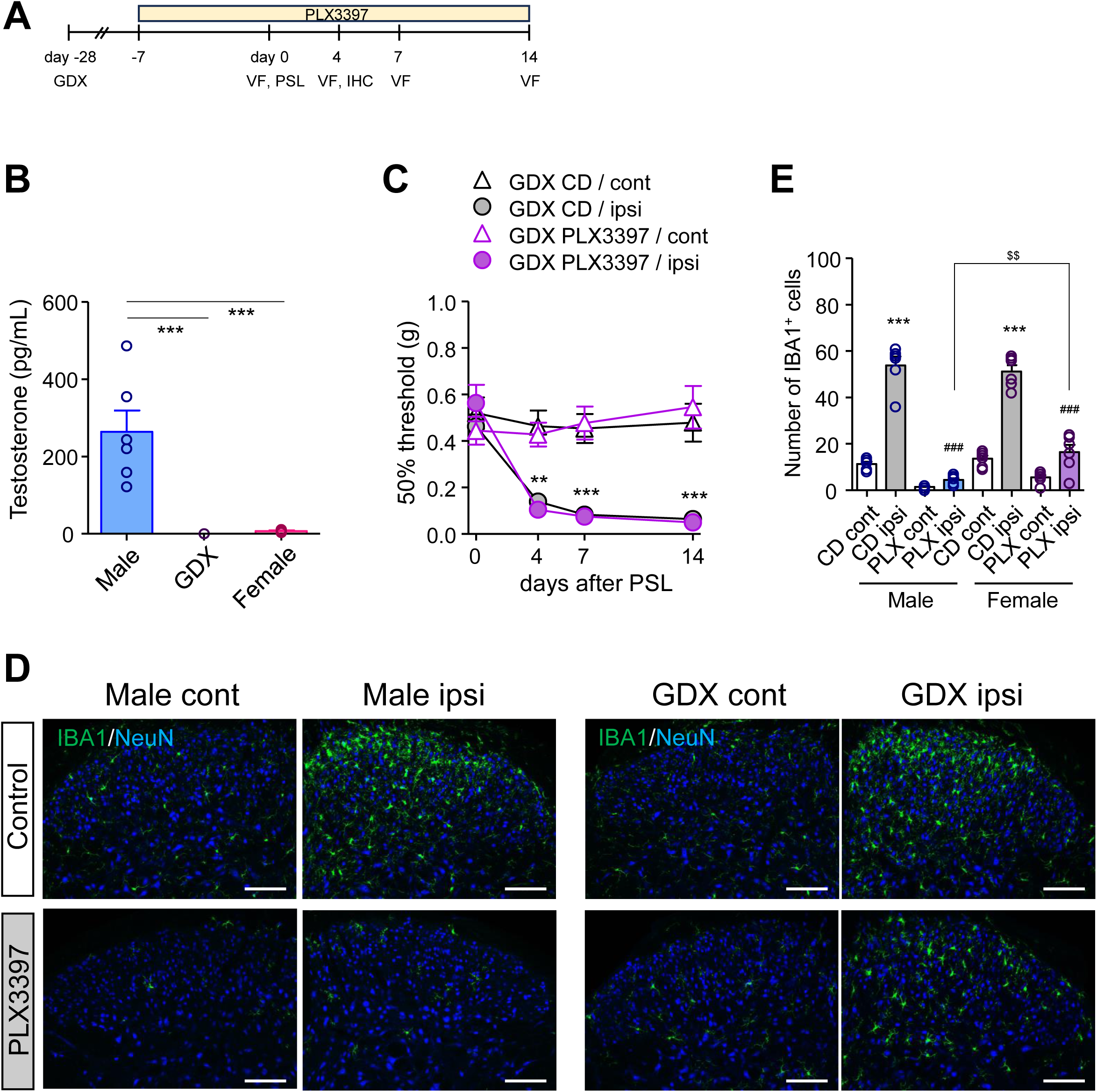
Androgen-dependent suppressive effects of PLX3397 on mechanical allodynia and microglial activation in SDH following PSL. Normal male and GDX male mice were fed a control diet (CD) or pexidartinib (PLX3397) diet and subsequently subjected to PSL. A) Time schedules of experiments. B) Serum testosterone levels of normal male, GDX male (four weeks post-surgery), and normal female mice were measured by LC-MS/MS. C) Effects of PLX3397 on the 50% mechanical threshold in normal male and GDX male mice were assessed by the up-down method using the von Frey test. D) Effects of PLX3397 on the expression of IBA1 in the SDH of normal male and GDX male mice were analyzed by IHC. Scale bars=100 μm. E) Quantitative analyses of the number of IBA1^+^ cells within the lamina I-III in the SDH on day 4 after PSL. Data are presented as the mean ± standard error of the mean (SEM); n=5-6. \*\*\**P*<0.001, \*\**P*<0.01 versus CD cont. ^###^*P*<0.001 versus CD ipsi. ^$$^*P*<0.01 versus PLX ipsi. GDX, gonadectomized; IHC, immunohistochemistry; LC-MS/MS, liquid chromatography-tandem mass spectrometry; PSL, partial sciatic nerve ligation; SDH, spinal dorsal horn.

To confirm the suppressive effects of PLX3397 on spinal microglial activation, we evaluated mRNA expression levels of microglial markers and inflammatory mediators in the SDH using RT-qPCR. Treatment with PLX3397 could markedly attenuate PSL-induced upregulation of IBA1, CD68, CX3CR1, CSF1R, and CCR5 in the SDH of normal male, GDX male, and female mice. Nevertheless, the degree of reduction in these molecules was greater in normal male mice than in GDX male or female mice, indicating that microglia in GDX male and female mice were less susceptible to the effects of PLX3397. Interestingly, treatment with PLX3397 after PSL significantly reduced the expression level of CCL3 in the SDH of male, but not GDX or female, mice. Moreover, the expression level of IL-1β showed a tendency for reduction in male, but not GDX or female, mice (**Fig. 4**).

**Figure 4.**
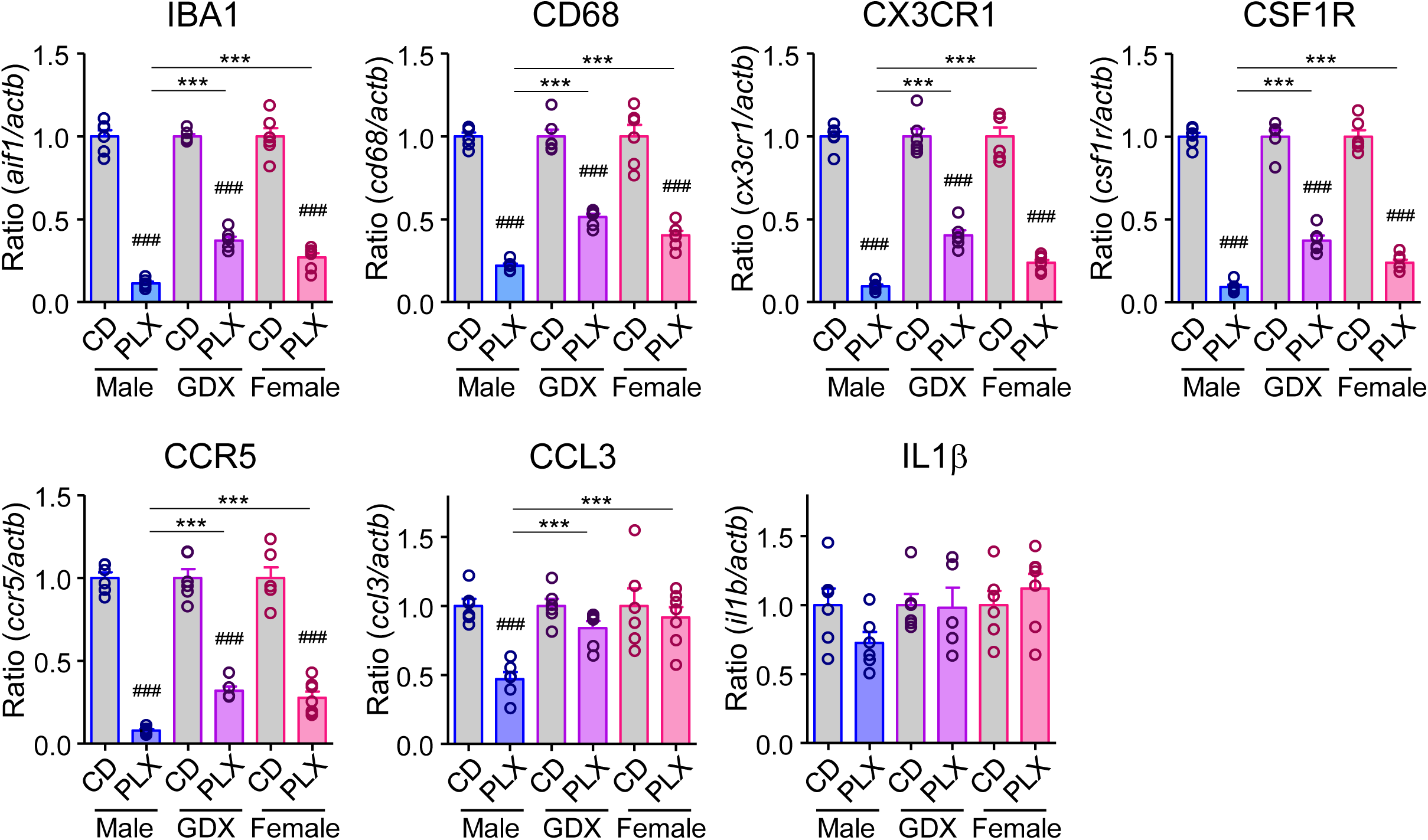
Sex- and androgen-dependent suppressive PLX3397 effects on microglial marker and inflammatory mediator upregulation in SDH. Male, GDX male, and female mice were fed a control diet (CD) or pexidartinib (PLX3397) diet, followed by PSL. The lumbar ipsilateral SDH was collected on day 7 after PSL. mRNA expression levels of *aif1*, *cd68*, *cx3cr1*, *csf1r*, *ccr5*, *ccl3*, and *il1b* were analyzed by RT-qPCR. Data are presented as the mean ± standard error of the mean (SEM); n=5-7. ^###^*P*<0.001 versus CD. \*\*\**P*<0.001 versus male PLX. GDX, gonadectomized; PSL, partial sciatic nerve ligation; SDH, spinal dorsal horn.

### Androgen-dependent suppressive effects of PLX3397 on peripheral macrophage activation and inflammatory mediators in the DRG following PSL

We next evaluated potential sex differences in mRNA expression levels of macrophage markers (IBA1, F4/80, CX3CR1, and CSF1R) and IL-1β, CCL3, and CCR5 in the DRG following PSL. All macrophage markers and CCR5 were similarly upregulated in both male and female mice. Although mRNA expression levels of these genes were slightly higher in females than in males, no significant sex-related differences were observed. Conversely, CCL3 and IL-1β showed sex-dependent mRNA expression patterns on indicated days (**Fig. 5**).

**Figure 5.**
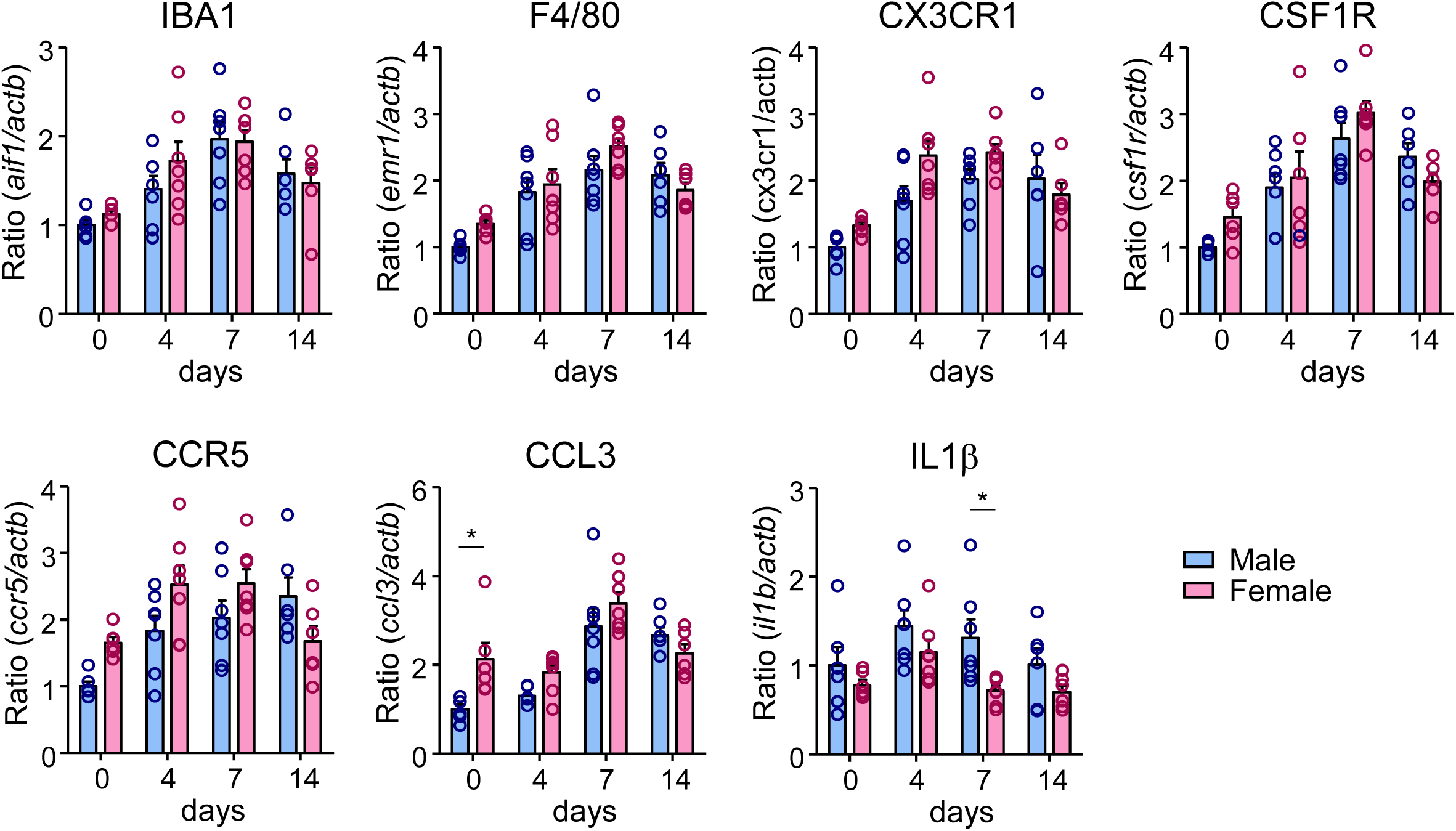
Upregulation of macrophage markers and inflammatory mediators in the DRG following PSL. Male and female mice were subjected to PSL, and the lumbar DRG were collected. mRNA expression levels of *aif1*, *emr1*, *cx3cr1*, *csf1r*, *ccr5*, *ccl3*, and *il1b* on days 0 (pre), 4, 7, and 14 after PSL were analyzed by RT- qPCR. Data are presented as the mean ± standard error of the mean (SEM); n=6-7. \**P*<0.05 versus male. DRG, dorsal root ganglia; PSL, partial sciatic nerve ligation.

Furthermore, we evaluated the suppressive effects of PLX3397 on the increased number of macrophages in the DRG following PSL induction. Based on IHC analysis, IBA1^+^ macrophages were markedly increased on the ipsilateral side of the DRG on day 7 post-PSL compared with those on the contralateral side in normal male and GDX male and female mice. Treatment with PLX3397 markedly reduced the number of IBA1^+^ macrophages on the ipsilateral side in all groups. Similar to spinal microglia, the number of remaining IBA1^+^ macrophages in GDX mice was slightly, but significantly, greater than that in male mice (**Fig. 6A, B**).

**Figure 6.**
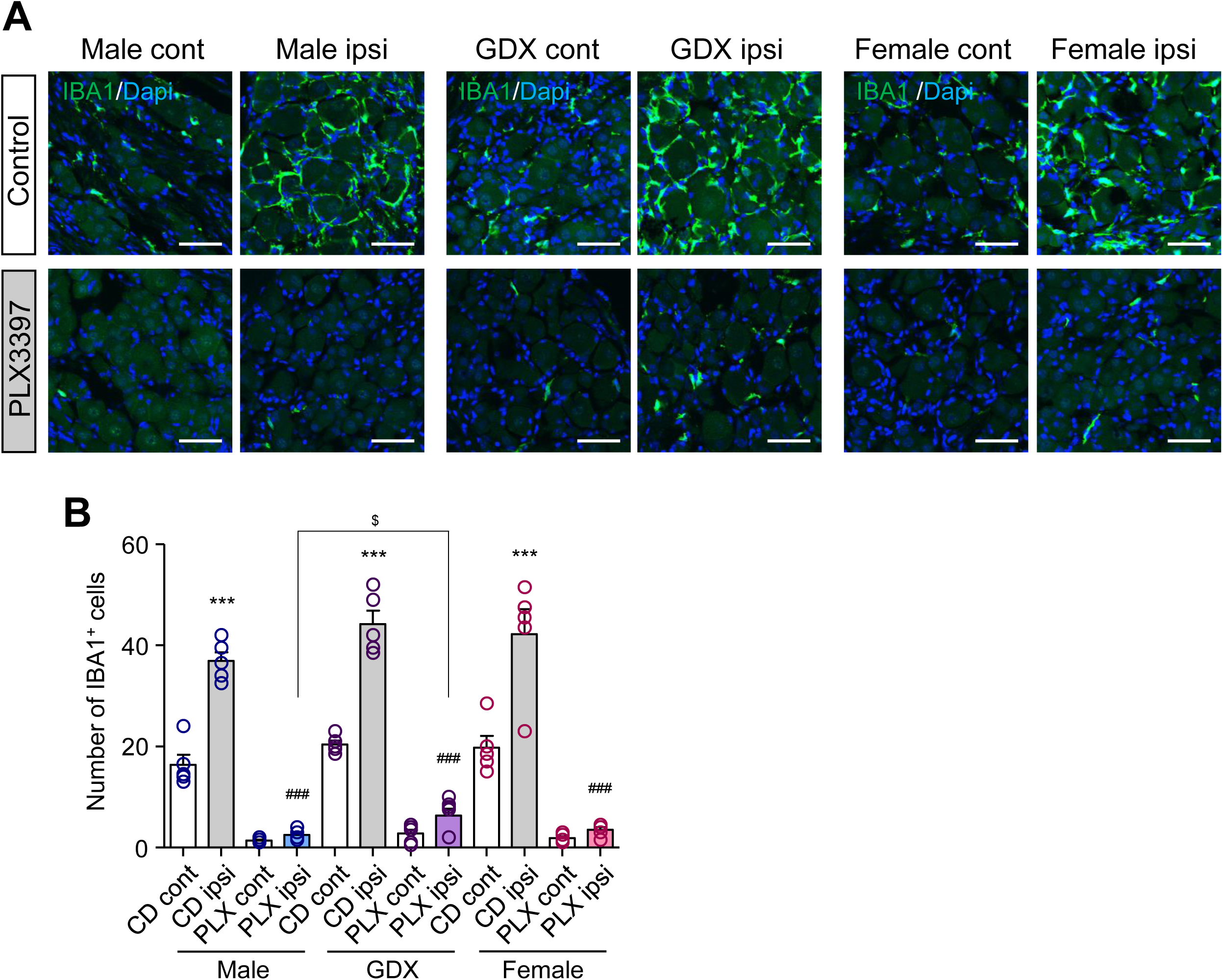
Sex- and androgen-dependent suppressive effects of PLX3397 on macrophage accumulation in DRG following PSL. Male, GDX male, and female mice were fed a control diet (CD) or pexidartinib (PLX3397) diet and subjected to PSL. The lumbar DRG was collected on day 7 after PSL. A) IBA1^+^ macrophages in the DRG were visualized by IHC. Scale bars=50 μm. B) Quantitative analyses of the number of IBA1^+^ cells within a square of 200 × 200 μm^2^ area. Data are presented as the mean ± standard error of the mean (SEM); n=5-6. \*\*\**P*<0.001 versus CD cont. ^###^*P*<0.001 versus CD ipsi. ^$^*P*<0.05 versus PLX ipsi. DRG, dorsal root ganglia; GDX, gonadectomized; IHC, immunohistochemistry; PSL, partial sciatic nerve ligation.

Consistent with the IHC results, we observed that treatment with PLX3397 markedly attenuated mRNA expression levels of IBA1, F4/80, CX3CR1, CSF1R, and CCR5 in the DRG of all mouse groups. However, the degree of reduction in these molecules was greater in normal males than in GDX male or female mice, indicating that DRG macrophages also showed androgen-dependent susceptibility to PLX3397. Although PLX3397 treatment significantly reduced expression levels of CCL3 and IL-1β expression, no sex- or androgen-dependent differences were observed (**Fig. 7**).

**Figure 7.**
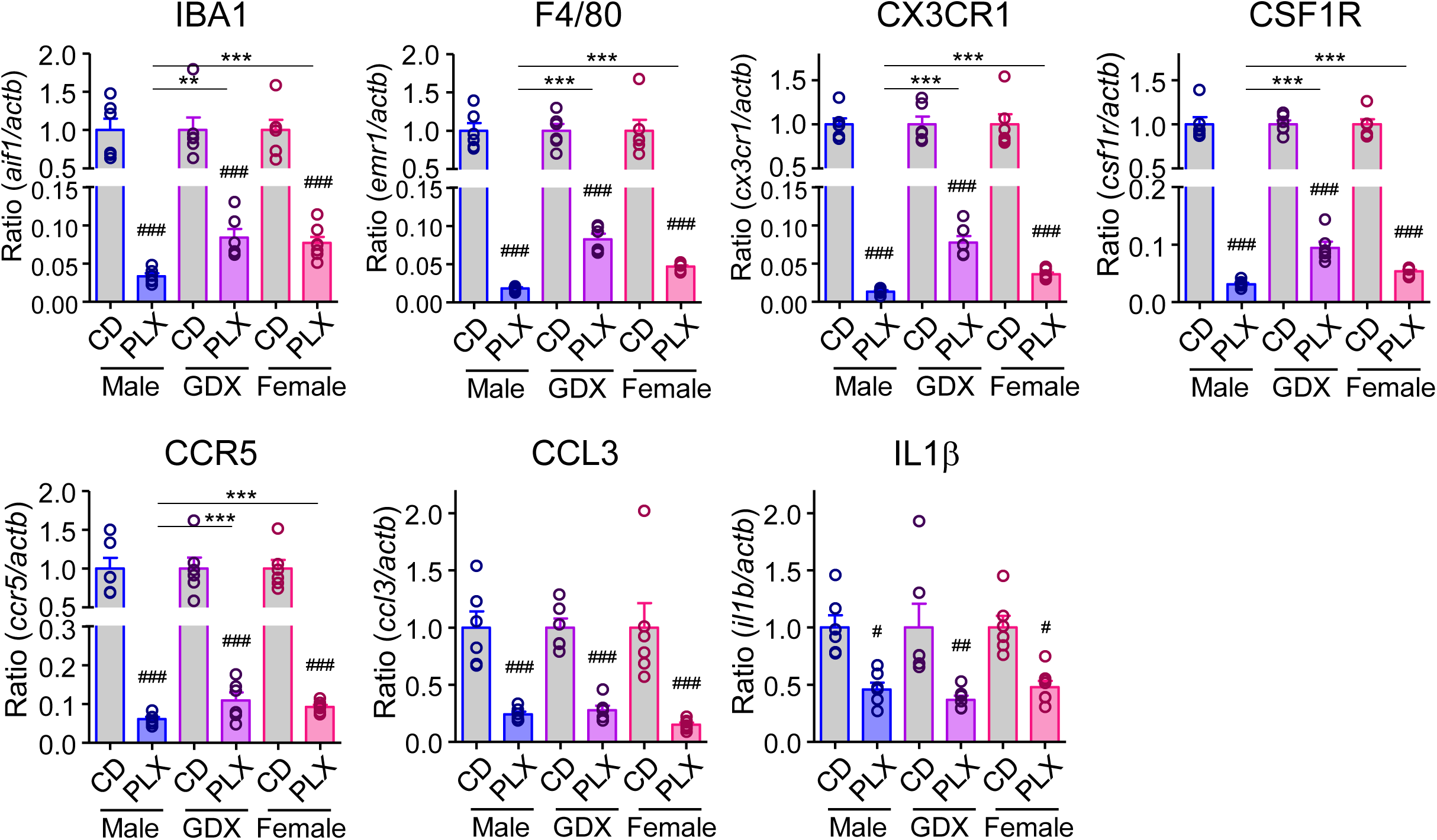
Sex- and androgen-dependent suppressive PLX3397 effects on macrophage marker and inflammatory mediator upregulation in DRG. Male, GDX male, and female mice were fed a control diet (CD) or pexidartinib (PLX3397) diet and subjected to PSL. The lumbar ipsilateral DRG were collected on day 7 after PSL. mRNA expression levels of *aif1*, *emr1*, *cx3cr1*, *csf1r*, *ccr5*, *ccl3*, and *il1b* were analyzed by RT-qPCR. Data are presented as the mean ± standard error of the mean (SEM); n=6-7. ^###^*P*<0.001, ^##^*P*<0.01, ^#^*P*<0.05 versus CD. \*\*\**P*<0.001, \*\**P*<0.01 versus Male PLX. DRG, dorsal root ganglia; GDX, gonadectomized; PSL, partial sciatic nerve ligation; RT-qPCR, reverse transcription-quantitative PCR.

### Effects of PLX3397 on peripheral macrophage activation and inflammatory mediators in the SCN following PSL

It is well-documented that the accumulation of inflammatory macrophages in the injured SCN plays a fundamental role in the development of neuropathic pain [5, 6, 10]. Therefore, we assessed whether sex- or androgen-dependent differences in the effects of PLX3397 can be observed in the SCN. Based on IHC analysis, F4/80^+^ macrophages were markedly increased on the ipsilateral side of the SCN on day 7 post-PSL when compared with those on the contralateral side in normal male and GDX male and female mice. Treatment with PLX3397 dramatically reduced the number of F4/80^+^ macrophages on the ipsilateral side in all mouse groups, with no significant differences observed in the suppressive effects of PLX3397 between these groups (**Fig. 8A, B**).

**Figure 8.**
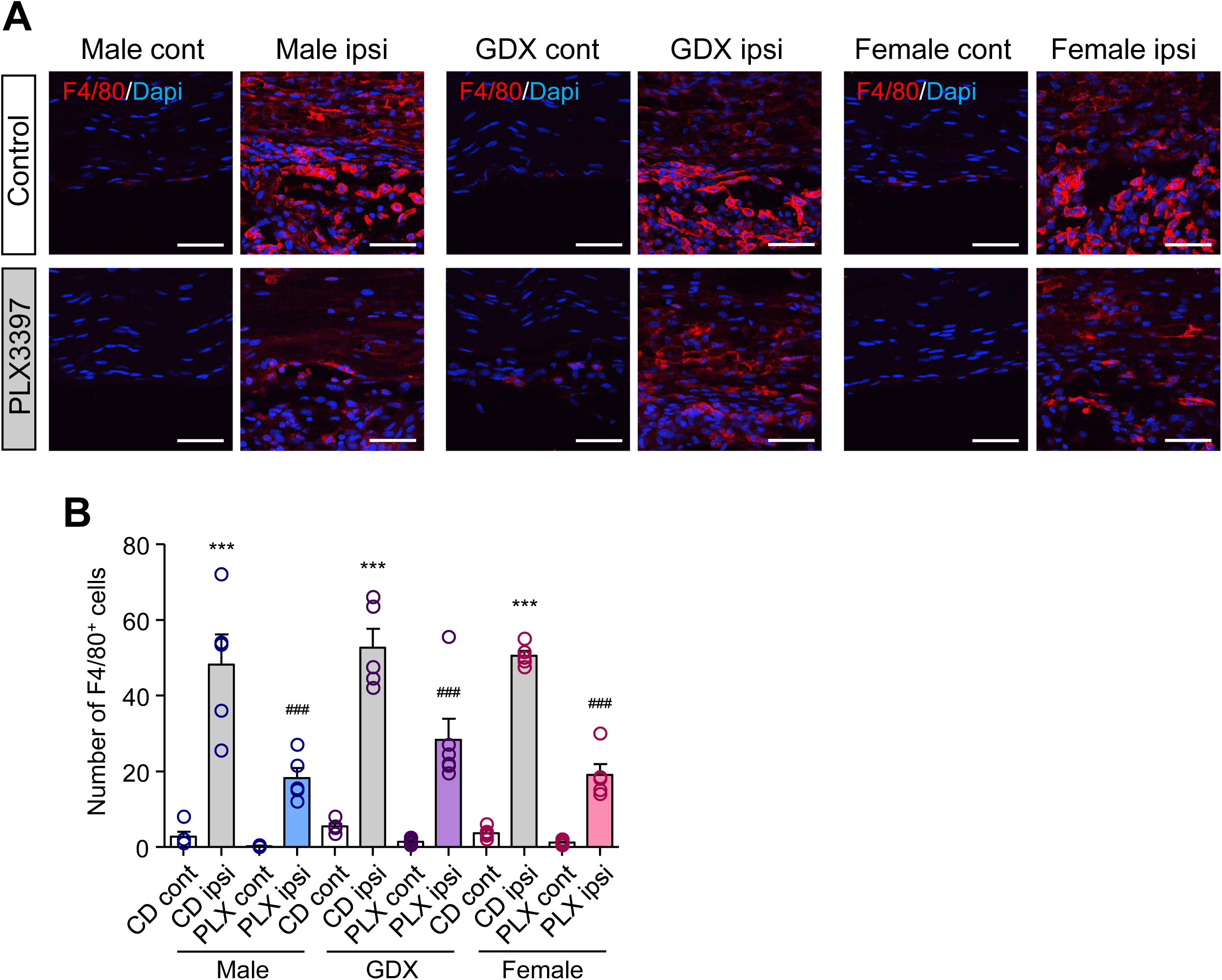
Suppressive effects of PLX3397 on macrophage accumulation in injured SCN following PSL. Male, GDX male, and female mice were fed a control diet (CD) or pexidartinib (PLX3397) diet and subjected to PSL. The SCN was collected on day 7 after PSL. A) F4/80^+^ macrophages in the SCN were visualized by IHC. Scale bars=50 μm. B) Quantitative analyses of the number of F4/80^+^ cells within a square of 200 × 200 μm^2^ area. Data are presented as the mean ± standard error of the mean (SEM); n=5-6. \*\*\**P*<0.001 versus CD cont. ^###^*P*<0.001 versus CD ipsi. IHC, immunohistochemistry; GDX, gonadectomized; PSL, partial sciatic nerve ligation; SCN, sciatic nerve.

The effects of PLX3397 on SCN macrophages were confirmed by detecting mRNA expression levels of IBA1, F4/80, CX3CR1, CSF1R, and CCR5 in the DRG after PSL induction, revealing that treatment with PLX3397 could similarly attenuate expression levels of these molecules in all experimental mouse groups. Unlike DRG, treatment with PLX3397 did not reduce mRNA expression levels of CCL3 and IL-1β in the SCN (**Fig. 9**). Considering the differences in the PLX3397-mediated suppressive effects on macrophage markers and inflammatory mediators between the DRG and SCN macrophages (**Fig. 8, 9**), the peripheral macrophages exhibited region specificity.

**Figure 9.**
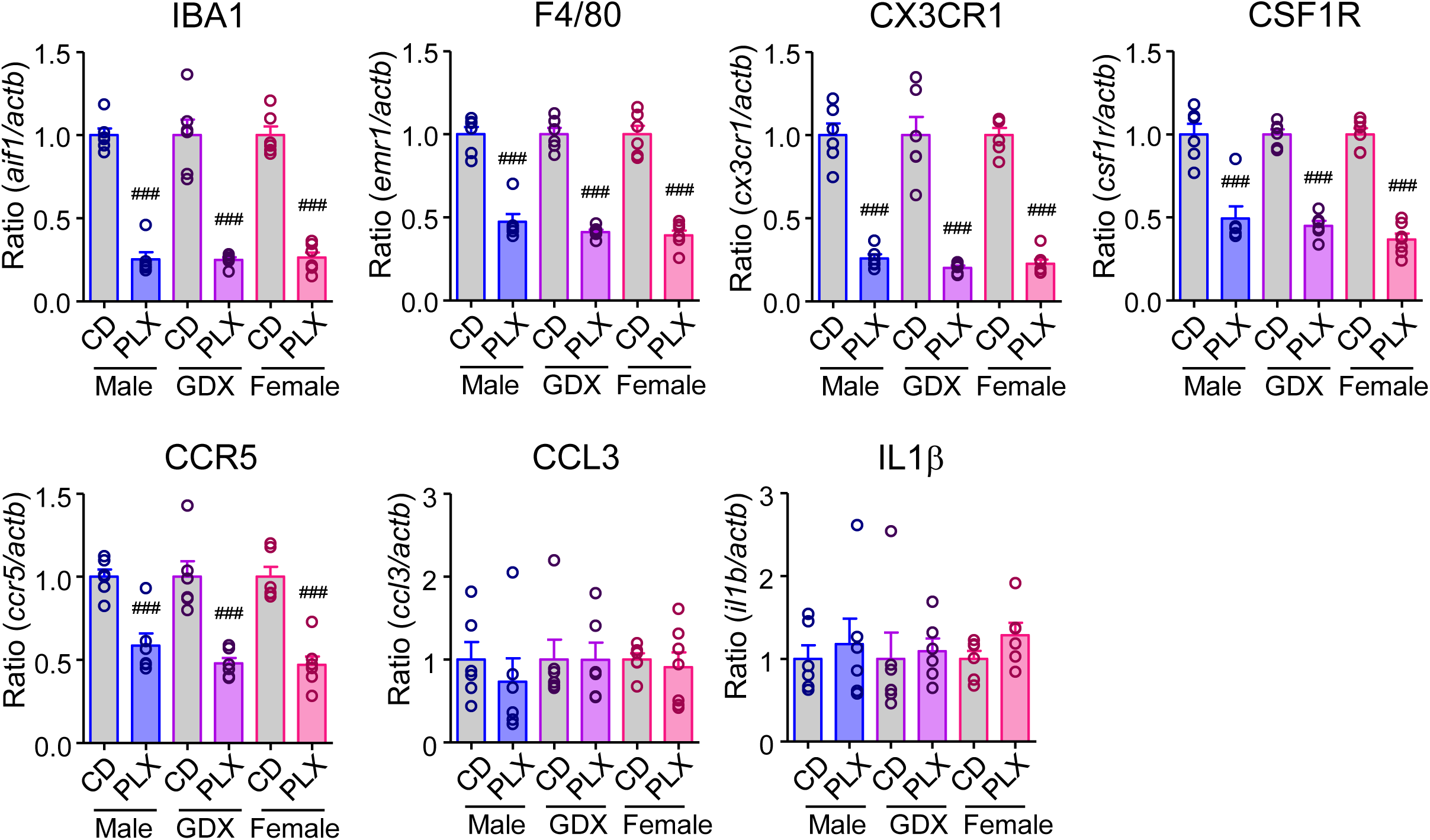
Suppressive effects of PLX3397 on macrophage marker and inflammatory mediator upregulation in injured SCN post-PSL. Male, GDX male, and female mice were fed a control diet (CD) or pexidartinib (PLX3397) diet and subjected to PSL. The injured SCN was collected on day 7 after PSL. The mRNA expression levels of *aif1*, *emr1*, *cx3cr1*, *csf1r*, *ccr5*, *ccl3*, and *il1b* were analyzed by RT-qPCR. Data are presented as the mean ± standard error of the mean (SEM); n=6-7. ^###^*P*<0.001 versus CD. GDX, gonadectomized; PSL, partial sciatic nerve ligation; RT-qPCR, reverse transcription-quantitative PCR; SCN, sciatic nerve.

CCL3 and IL-1β were found to be critical mediators underlying the etiology of neuropathic pain [9, 33]. Given the absence of sex- or androgen-dependent differences in SCN macrophages, we hypothesized that blockade of CCR5, a principal receptor of CCL3 in the injured SCN, might prevent neuropathic pain in both sexes. As expected, perineural administration of maraviroc, a CCR5 antagonist, significantly prevented PSL-induced mechanical allodynia in both male and female mice (**Fig. 10**). These results suggest that inflammatory mechanisms associated with the SCN are commonly observed in both sexes and contribute to the etiology of neuropathic pain.

**Figure 10.**
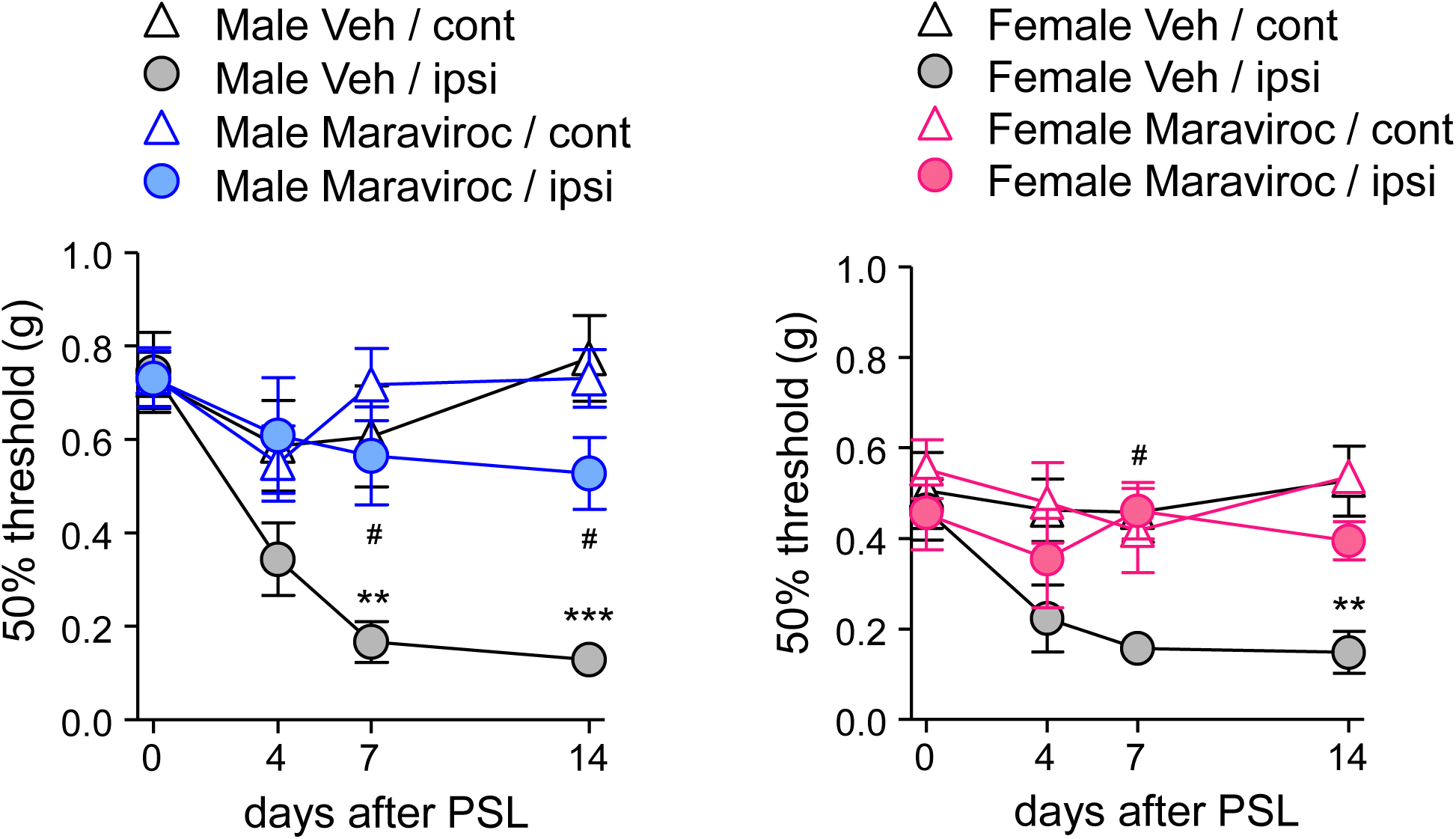
Prevention of mechanical allodynia by inhibiting the CCL3-CCR5 system in both sexes. Male and female mice were subjected to PSL, and maraviroc (20 nmol) was perineurally administered for 7 consecutive days (days 0–6) after PSL. The effects of maraviroc on the 50% mechanical threshold in male and female mice were assessed with the up-down method using the von Frey test. Data are presented as the mean ± standard error of the mean (SEM); n=6. \*\*\**P*<0.001, \*\**P*<0.01 versus Veh/cont. ^#^*P*<0.05 versus Veh/ipsi. PSL, partial sciatic nerve ligation.

## Discussion

In the current study, we highlighted the sex- and androgen-dependent characteristics of spinal microglia and peripheral macrophages that underlie the etiology of neuropathic pain. Both male and female mice exhibited marked microglial activation in the SDH and peripheral macrophage accumulation in the DRG and SCN after PSL. PLX3397-mediated blockade of CSF1R substantially suppressed PSL-induced mechanical allodynia in male but not in female mice. Although activated microglia in the SDH and inflammatory macrophages in the DRG were markedly decreased in both male and female mice, there were certain sex differences in the PLX3397-induced depletive effects on microglia and macrophages between male and female mice, that is, the number of remaining microglia and DRG macrophages in female was greater than that in male mice. Importantly, male GDX mice showed similar results as female mice, suggesting that intrinsic androgen levels might determine the susceptibility of microglia and macrophages to the effects of PLX3397. In contrast to the effects on DRG, treatment with PLX3397 did not exhibit sex-dependent differences in the SCN macrophages, and inhibition of CCL3-CCR5 signaling by the local administration of maraviroc successfully prevented PSL-induced mechanical allodynia in both male and female mice.

CSF1R, expressed on macrophages and microglia, functions as a key regulator of myeloid cell proliferation, differentiation, and survival [28, 29]. It was recently reported that the upregulation and release of CSF1 from primary sensory neurons following nerve injury can subsequently lead to the proliferation and activation of spinal microglia, contributing to neuropathic pain [14]. Moreover, CSF1 is reportedly released from the cell body of sensory neurons and also plays a critical role in macrophage proliferation and accumulation in the DRG [7, 37]. Therefore, we expected that CSF1R antagonists would inhibit and deplete not only spinal microglia but also peripheral macrophages. Among several inhibitors, PLX3397 is orally active and extensively employed to investigate the pathophysiological roles of microglia and macrophages in various neuroinflammation-associated diseases [38–40]. In the field of pain, PLX3397 (or PLX5622) has been used to determine the role of microglia and macrophages in different pain modalities. In addition to acute inflammatory and bone cancer pain, treatment with PLX3397 was shown to substantially suppress chronic pain caused by limb ischemia or nerve injury [41–43]. However, it is worth noting that the majority of the evidence is based on findings from male animals without analyzing sex differences. To the best of our knowledge, we are the first to demonstrate the sex- and androgen-dependent suppressive effects of PLX3397 on neuropathic pain.

To date, several key components that define reactive microglia, such as interferon regulatory factor 5 (IRF5) and IRF8, have been identified [16, 44]. Although it remains unclear whether these factors exert sex-dependent actions, the purinergic P2X4 receptor (P2X4R) and p38-mitogen-activated protein kinases (p38 MAPK) are known to be predominant in males, contributing to neuropathic pain in male but not female mice [27]. Similar to general microglial inhibitors such as minocycline, P2X4R or P38 MAPK inhibitors fail to prevent neuropathic pain in female mice [19, 20]. Based on these findings, characteristics of activated microglia may differ between male and female mice, even if morphological changes and expression levels of general microglial markers (i.e., IBA1, CX3CR1, and CSF1R) are similarly observed in both sexes, as supported by transcriptome analyses of microglia following nerve injury [22, 23]. Furthermore, manipulating spinal microglia by administering lipopolysaccharides or via chemogenetic approaches elicits pain hypersensitivity in male but not female mice with sex-dependent gene expression profiles [21, 26]. Herein, we also confirmed that PLX3397 treatment could suppress neuropathic pain only in normal male but not GDX male or female mice, despite depletion of the majority of microglia, indicating that the depletion of spinal microglia cannot suppress neuropathic pain in mice with low androgen levels.

In addition to spinal microglia, peripheral macrophages play a fundamental role in the etiology of neuropathic pain, given that myeloid cells in both the PNS and CNS synergistically accelerate neuroinflammation [31, 45]. Inflammatory macrophages have been recently identified in the DRG, as the depletion of DRG macrophages prevented neuropathic pain after nerve injury in both male and female mice. Following nerve injury, macrophage accumulation in the DRG was also based on CSF1 production in DRG neurons [37]. However, our results revealed that treatment with PLX3397 did not suppress neuropathic pain in female and GDX male mice, although PLX3397 depleted most DRG macrophages, indicating that the contribution of DRG macrophages to neuropathic pain is also male-dominant, similar to spinal microglia. One possible explanation for this discrepancy is the use of different nerve injury models in each study. In contrast, our group reported that inflammatory macrophages in the injured SCN are a principal component of neuropathic pain. Inhibition or depletion of SCN macrophages prevents nerve injury- and diabetes-associated neuropathic pain [46–48], and macrophages in peripheral nerves reportedly exert sex-independent pain-facilitating effects [49]. Importantly, numerous SCN macrophages could be observed even after PLX3397 treatment when compared with DRG macrophages. Our results suggest that the basic characteristics of macrophages differ between the DRG and SCN; that is, DRG macrophages mainly proliferate from resident cells, such as spinal microglia, while SCN macrophages are recruited from the bone marrow [50] and exhibit common functions in male and female mice.

Activated microglia and macrophages produce several inflammatory cytokines (e.g., IL-1β and tumor necrosis factor α) and chemokines that directly increase the excitability of pain-processing neurons in the PNS and the CNS [3, 4, 8]. Among these inflammatory mediators, IL-1β is the most notable, as inhibition of IL-1β is well-known to prevent pain hypersensitivity [34, 51, 52]. Considering the absence of significant sex difference in the effects of PLX3397 on expression levels of IL-1β in the SDH, DRG, and SCN, clarifying the sex-dependent suppressive effects of PLX3397 by focusing on IL-1β could present a considerable challenge. Along with other researchers, we have previously demonstrated that CCL3 plays a crucial role in the etiology of neuropathic pain. CCL3 is upregulated in the SDH after nerve injury, and its receptor CCR5 is mainly expressed on microglia, indicating that the CCL3-CCR5 system underlies neuropathic pain [53, 54]. Interestingly, treatment with PLX3397 partially, but significantly, reduced the expression level of CCL3 in male, but not female or GDX male, mice, suggesting sex- and androgen-dependent differences in the gene expression profiles of microglia. The pivotal role of CCL3 in neuropathic pain has been examined in the PNS [33, 55]. CCL3 is rapidly upregulated by nerve injury and facilitates neuropathic pain via the CCR5-mediated upregulation of IL-1β in the SCN [34, 56]. Given that CCL3 and IL-1β are expressed in both macrophages and Schwann cells, the fact that CCL3 expression levels were not reduced could be explained by remaining macrophages and Schwann cells functioning as primary sources of CCL3 in the injured SCN.

The relationship between sex hormones (i.e., androgens and estrogens) and the pathogenesis of inflammatory diseases has been discussed previously [57, 58]. Indeed, several reports have shown that the immune system, including microglia and macrophages, differs between males and females [59–61]. However, the mechanisms through which sex hormones determine sex-dominant characteristics of microglia and macrophages are poorly understood. Herein, we found that spinal microglia and DRG macrophages show clear sex differences in the etiology of neuropathic pain based on intrinsic androgen levels. GDX male mice showed myeloid cell phenotypes and serum androgen levels similar to those of female mice. However, evidence that microglia and macrophages express androgen receptors remains insufficient despite sex-related differences in myeloid cells in several diseases [58, 59]. Although we explored androgen receptor-expressing cells in the SDH and DRG, we did not observe the obvious expression of androgen receptors in either microglia or macrophages. Moreover, sex hormones may largely affect pain sensitivity [62, 63], given that androgens (testosterone) exert antinociceptive effects in several models of pathological pain [35, 36]. To clarify the mechanisms underlying sex differences in pain processing, it is imperative to establish androgen-targeted cells.

Collectively, the findings of the current study demonstrated substantial sex- and androgen-dependent differences in the etiology of neuropathic pain in spinal microglia and DRG macrophages. Notably, treatment with PLX3397 could successfully suppress neuropathic pain in male, but not female, mice, although PLX3397 drastically depleted activated spinal microglia and DRG macrophages that might have proliferated from resident cells in both sexes. We also found that androgen levels determine the sex-dependent roles of spinal microglia and DRG macrophages in pain processing. More importantly, SCN macrophages did not exhibit any sex-related differences in the etiology of neuropathic pain, and the CCL3-CCR5 system commonly contributed to neuropathic pain in male and female mice. Given that the mechanisms of neuropathic pain may differ among experimental models and clinical conditions, it is necessary to accumulate several lines of evidence to understand the sex-dependent regulatory mechanisms of pain.

## Abbreviations

CCL3: CC-chemokine ligand 3
CCR5: CC-chemokine receptor 5
CNS: central nervous systems
CX3CR1: CX3C-chemokine receptor 1
DRG: dorsal root ganglia
CSF1: colony-stimulating factor 1
CSF1R: CSF1 receptor
GDX: gonadectomized
IBA1: ionized calcium-binding adapter molecule 1
IHC: immunohistochemistry
IL: interleukin
IRF: interferon regulatory factor
PBS: phosphate-buffered saline
PNS: peripheral nervous system
PSL: partial sciatic nerve ligation
P2X4R: purinergic P2X4 receptor
p38 MAPK: p38-mitogen-activated protein kinases
RT-qPCR: reverse transcription-quantitative PCR
SCN: sciatic nerve
SDH: spinal dorsal horn

## Declarations

### Ethics approval and consent to participate

All animal experiments were approved by the Animal Research Committee of Wakayama Medical University and performed in accordance with the in-house guidelines for the care and use of laboratory animals of Wakayama Medical University and the Animal Research: Reporting of In Vivo Experiments (ARRIVE) guidelines.

### Consent for publication

Not applicable

### Availability of data and materials

The datasets used and/or analyzed in the current study are available from the corresponding author upon reasonable request.

### Author contributions

FS, YF, and NK conducted the experiments. YF, YH, SK, SH, and KS analyzed the data. NK supervised the study. FS and NK prepared the manuscript. All the authors have read and approved the final version of the manuscript.

## Acknowledgments

We want to thank Editage (https://www.editage.com) for the English language editing.

