## Supplementary information for "Sexually dimorphic effects of pexidartinib on nerve injury-induced neuropathic pain in mice"

**Table S1.** Primer sequences for reverse transcription-quantitative PCR (RT-qPCR).

**Figure S1**. Effects of PLX3397 on the expression of glial markers, pain sensitivity, and motor function. Male mice were fed a control diet (CD) or pexidartinib (PLX3397) diet, and the lumbar SDH was collected. A) mRNA expression levels of *aif1* at weeks 0 (pre), 1, 2, and 3 during treatment and *gfap* at 2 weeks of treatment were analyzed by RT-qPCR. Effects of PLX3397 on 50% mechanical threshold, thermal withdrawal latency, and motor function were measured using the von Frey test (B), Hargreaves test (C), and rotarod test (D), respectively, at 2 weeks of PLX3397 treatment. Data are presented as the mean ± standard error of the mean (SEM); n=5-6. ***$P$<0.001 versus 0 week (pre). PSL, partial sciatic nerve ligation; RT-qPCR, reverse transcription-quantitative PCR; SDH, spinal dorsal horn.

Table S1. Primer sequences for RT-qPCR.

| Gene | Forward (5' to 3') | Reverse (5' to 3') |
|---|---|---|
| *actb* | CAGCTGAGAGGGAAATCGTG | TCTCCAGGGAGGAAGAGGAT |
| *aif1* | ATGAGCCAAAGCAGGGATTT | TTGGGATCATCGAGGAATTG |
| *cd11b* | GTTTCTACTGTCCCCCAGCA | GTTGGAGCCGAACAAATAGC |
| *cd68* | ACTCATAACCCTGCCACCAC | CCAACAGTGGAGGATCTTGG |
| *cx3cr1* | CCTCCTTCCCTGAACTGGAT | GGACAGGAAGATGGTTCCAA |
| *csf1r* | TCTCCCTACTGGACCTTGGA | GGGTCTTCAAGCTCGGTACA |
| *ccr5* | CGAAAACACATGGTCAAACG | TCTCCTGTGGATCGGGTATA |
| *ccl3* | CTGCCCTTGCTGTTCTTCTC | GTGGAATCTTCCGGCTGTAG |
| *il1b* | AAAGCTCTCCACCTCAATGG | AGGCCACAGGTATTTTGTCG |
| *emr1* | AACTTTCAAGGCCCAGGAGT | GCTCTCCCCAGGATATTGGT |

**Figure S1**

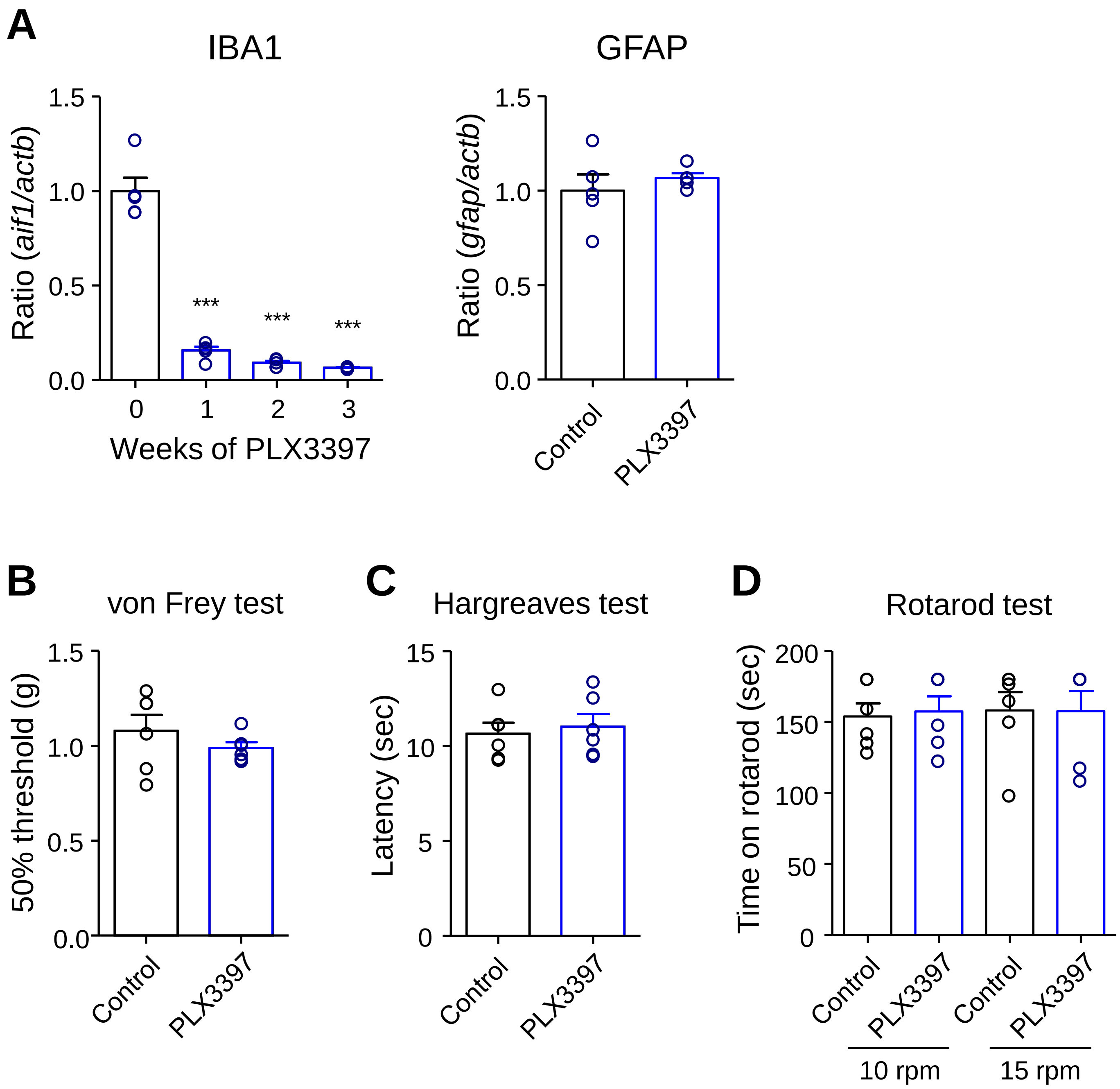
A
IBA1
Ratio (*aif1/actb*)
1.5
1.0
0.5
0.0
***
***
***
0
1
2
3
Weeks of PLX3397
GFAP
Ratio (*gfap/actb*)
1.5
1.0
0.5
0.0
Control
PLX3397
B
von Frey test
50% threshold (g)
1.5
1.0
0.5
0.0
Control
PLX3397
C
Hargreaves test
Latency (sec)
15
10
5
0
Control
PLX3397
D
Rotarod test
Time on rotarod (sec)
200
150
100
50
0
Control
PLX3397
Control
PLX3397
10 rpm
15 rpm
